# Glutamate uptake is transiently compromised in the perilesional cortex following controlled cortical impact

**DOI:** 10.1101/2024.08.28.610143

**Authors:** Jacqueline P Garcia, Moritz Armbruster, Mary Sommer, Aliana Nunez-Beringer, Chris G Dulla

## Abstract

Glutamate, the primary excitatory neurotransmitter in the CNS, is regulated by the excitatory amino acid transporters (EAATs) GLT-1 and GLAST. Following traumatic brain injury (TBI), extracellular glutamate levels increase, contributing to excitotoxicity, circuit dysfunction, and morbidity. Increased neuronal glutamate release and compromised astrocyte-mediated uptake contribute to elevated glutamate, but the mechanistic and spatiotemporal underpinnings of these changes are not well established. Using the controlled cortical impact (CCI) model of TBI and iGluSnFR glutamate imaging, we quantified extracellular glutamate dynamics after injury. Three days post-injury, glutamate release was increased, and glutamate uptake and GLT-1 expression were reduced. 7- and 14-days post-injury, glutamate dynamics were comparable between sham and CCI animals. Changes in peak glutamate response were unique to specific cortical layers and proximity to injury. This was likely driven by increases in glutamate release, which was spatially heterogenous, rather than reduced uptake, which was spatially uniform. The astrocyte K^+^ channel, Kir4.1, regulates activity-dependent slowing of glutamate uptake. Surprisingly, Kir4.1 was unchanged after CCI and accordingly, activity-dependent slowing of glutamate uptake was unaltered. This dynamic glutamate dysregulation after TBI underscores a brief period in which disrupted glutamate uptake may contribute to dysfunction and highlights a potential therapeutic window to restore glutamate homeostasis.

## INTRODUCTION

Traumatic brain injury (TBI) affects more than 60 million people worldwide (Dewan et al. 2018) and sets into motion a complex series of molecular, cellular, circuit, and systems-level events that promote functional recovery, but can also drive secondary injury (Lutkenhoff et al. 2020). Multiple studies show that levels of glutamate, the primary excitatory neurotransmitter in the central nervous system (Bergles et al. 1999; Clements et al. 1992; Diamond and Jahr 1997; Marvin et al. 2013), are increased in the human brain (Katayama et al. 1990) and in animal models (Sowers et al. 2021) after TBI. In patients, increases in glutamate occur early after injury, persist for days, and are associated with increased mortality (Chamoun et al. 2010). In animals, similar changes occur, but resolve more rapidly (Sowers et al. 2021). These studies use *in vivo* micro-dialysis which allows for quantitative analysis of glutamate concentrations but has poor spatial and temporal resolution. Increases in glutamate release after TBI likely occur due to acute injury-induced cell rupture (Qin et al. 2021), enhanced neuronal release of glutamate (Hinzman et al. 2010), and disrupted uptake of glutamate by astrocytes (Sowers et al. 2021). How these dynamically contribute to injury-induced elevations in glutamate after TBI is not well established, but multiple studies implicate the loss of the astrocyte excitatory amino acid transporters (EAATs) GLT-1 and GLAST (Ventura and Harris 1999) both in animal models (David et al. 2009; Shandra et al. 2019; van Landeghem et al. 2006) and in patients (Beschorner et al. 2007; van Landeghem et al. 2006). Loss of EAATs would compromise glutamate clearance, potentially contributing to synaptic and circuit dysfunction. Restoring normal glutamate levels after TBI could prevent pathological excitation, excitotoxicity, and improve patient outcomes after TBI. Thus, it is critical to understand the mechanisms leading to elevated glutamate levels after TBI.

Astrocytes control neuronal function by modulating synaptic transmission, ion gradients, and more, and they sense and rapidly respond to synaptic activity (Anderson and Swanson 2000; Araque et al. 1998). Astrocytes undergo structural, functional, and molecular changes following TBI (Latov et al. 1979; Palmer et al. 1993; Susarla et al. 2014; Takamiya et al. 1988; Villapol et al. 2014), but how injury affects glutamate clearance, and the activity-dependent slowing of glutamate uptake is not fully understood (Armbruster et al. 2016; Armbruster et al. 2022). When glutamate is released by neurons, it is rapidly bound by GLT-1 and GLAST (Ventura and Harris 1999) and transported into astrocytes (although neurons also express GLT-1 (Chen et al. 2004a)). This ensures the spatiotemporal precision of synaptic transmission and prevents elevations of extracellular glutamate (Ventura and Harris 1999). Transport of glutamate via EAATs are both sodium-driven and voltage-dependent (Barbour et al. 1988; Bergles et al. 2002; Kanai et al. 1994; Klockner et al. 1993; Levy et al. 1998; Longuemare et al. 1999; Zhou et al. 2014). GLT-1 is the predominant EAAT in the adult brain (Danbolt 2001), and even modest reductions in GLT-1 impair glutamate clearance. Astrocytes also buffer extracellular K^+^, via the inwardly rectifying K^+^ channel Kir4.1, to maintain ion gradients essential to controlling neuronal and glial function (Higashi et al. 2001). We recently showed that neuronal activity can rapidly inhibit glutamate uptake with synapse specificity, via local depolarization of peripheral astrocyte processes (Armbruster et al. 2016; Armbruster et al. 2022). Because EAAT function is voltage-dependent (Bergles et al. 2002; Levy et al. 1998) and increases in extracellular K^+^ levels depolarize astrocyte membrane potential (V_m_) (Armbruster et al. 2022; Dezsi et al. 2016; Ransom and Goldring 1973), EAATs and Kir4.1 are thought to work together to maintain efficient and dynamic glutamate uptake (Barbour et al. 1988).

In this study, we used intensity-based glutamate sensing fluorescent reporter (iGluSnFR) (Marvin et al. 2019) to assay glutamate dynamics in the extracellular space following TBI. iGluSnFR is an extracellular membrane-tethered glutamate sensor that enables optical detection of synaptic glutamate release and clearance dynamics. iGluSnFR allows quantification of glutamate dynamics across large regions of the brain with high spatial and temporal resolution. Following TBI, glutamate release is enhanced (Folkersma et al. 2011; Katayama et al. 1990) and extracellular glutamate levels are elevated in patients (Lutkenhoff et al. 2020) and mouse models of TBI (Cantu et al. 2015; Nilsson et al. 1994). We do not understand, however, how TBI causes spatially-specific changes in glutamate release or glutamate uptake. iGluSnFR imaging allows a completely novel approach to define the spatiotemporal changes in glutamate signals that follow TBI. To model TBI, we used the controlled cortical impact (CCI) model, an established rodent model (Lighthall 1988; Lighthall et al. 1990; Osier and Dixon 2016) that induces cortical contusion, blood-brain-barrier (BBB) disruption, brain volume loss, secondary injury (apoptosis, inflammation, oxidative stress), ventricular enlargement, astrogliosis, cortical hyperexcitability, aberrant glutamate signaling, metabolic disruption, and learning, memory, motor, and executive dysfunction (Budinich et al. 2013; Song et al. 2016; Zhao et al. 2015).

Using the CCI model, we found that stimulus-evoked glutamate release was significantly increased, and glutamate uptake was significantly decreased in acute cortical brain slices 3 days after injury. 7 and 14-days after injury, glutamate dynamics were comparable between CCI and sham mice. Activity-dependent slowing of glutamate uptake was similar in the healthy and injured brain at all time points. Immunohistochemistry showed significantly decreased levels of GLT-1 expression 3 days after TBI, while GLAST and Kir4.1 expression was unchanged. GLT-1 expression levels returned to baseline 7 and 14-days after injury. Three days post-injury, spatial iGluSnFR analysis revealed that glutamate release was increased in superficial cortical layers, relative to deep layers as well as proximal to the site of injury versus distally, while the rate of glutamate clearance was comparable across regions. Our study shows that transient decreases in EAATs broadly compromise glutamate uptake after CCI, while changes in glutamate release are more focal, with no effect on activity-dependent inhibition of glutamate uptake. This suggests glutamate dysregulation may contribute to TBI related pathology in the days following injury and glutamate-based therapeutic approaches may be most effective when delivered during that time window.

## METHODS

### Adeno-associated virus injection

C57BL/6 male mice (35-45 days old) were stereotaxtically injected with AAV.php.EB-hSyn-SF-iGluSnFR.A184V (Virovek) (Marvin et al. 2018) in a single hemisphere with 3 injections sites (coordinates): (-1.25, -1.25, -0.5), (1.25, 2.25, 0.5), and (1.25, 3.25, 0.5) ( +x, +y, -z) mm as previously described (Armbruster et al. 2016; Armbruster et al. 2022). Mice were anesthetized with isoflurane for surgery, and reporter viruses were injected with 1 μl per site (0.1– 0.2 μl/min) at 4.5E9 gene copies. Mice were used for immunofluorescence or acute slice preparations 14– 28 d after injection. Mice were housed in 12 hour /12 hour light/dark cycles following surgeries. Procedures followed all guidelines of the Tufts University School of Medicine’s Institutional Animal Care and Use Committee.

### Controlled cortical impact model of TBI

We utilized a controlled cortical impact (CCI) model of traumatic brain injury, as previously described (Armbruster et al. 2022; Bottom-Tanzer et al. 2024; Cantu et al. 2015; Koenig et al. 2019). C57BL/6 mice (45-52 days old) underwent CCI or sham surgery. For CCI surgery, mice were anesthetized with isoflurane in O_2_ (4% induction, 2% maintenance) and placed in a Kopf stereotaxic frame. Following a midline incision, a 5-mm craniotomy was performed over the left sensorimotor cortex, lateral to the sagittal suture between bregma and lambda, ensuring no damage to the underlying dura. The craniotomy site was cooled via constant irrigation with sterile saline during drilling. The cortical lesion was induced with an electromagnetic impactor (Leica) using CCI parameters consistent with a moderate injury: 3-mm-diameter impactor tip, speed of 3.5 m/s, depth of 1 mm, and dwell time of 400ms. Excess bleeding was flushed away with sterile, room temperature saline prior to suturing. For sham surgeries, mice were anesthetized with isoflurane in O_2_ (4% induction, 2% maintenance) and placed in a Kopf stereotaxic frame for a time equivalent to the duration of a CCI surgery.

### Preparation of acute brain slices

400 μm thick cortical brain slices were prepared from CCI or sham mice expressing iGluSnFr (Armbruster et al. 2016). Mice were anesthetized with isoflurane, decapitated, and the brains were rapidly removed and placed in ice-cold slicing solution equilibrated with 95% O_2_:5% CO_2_ (in mM) as follows: 2.5 KCl, 1.25 NaH_2_PO_4_, 10 MgSO_4_, 0.5 CaCl_2_, 11 glucose, 234 sucrose, and 26 NaHCO_3_. The brain was glued to a Vibratome VT1200S (Leica), and slices were cut in a coronal orientation. Slices were then placed into a recovery chamber containing aCSF (in mM) as follows: 126 NaCl, 2.5 KCl, 1.25 NaH_2_PO_4_, 1 MgSO_4_, 2 CaCl_2_, 10 glucose, and 26 NaHCO_3_ (equilibrated with 95% O_2_:5% CO_2_) then equilibrated in aCSF at 32°C for 1 h. Slices were allowed to return to room temperature and used for live imaging.

### Live imaging

Glutamate imaging was performed as previously described (Armbruster et al. 2016; Armbruster et al. 2022). Briefly, brain slices expressing iGluSnFR were placed into a submersion chamber (Siskiyou), held in place with small gold wires, and perfused with aCSF containing 20 µm DNQX (antagonist of AMPA receptors), and 50 µm (APV, antagonist of NMDA receptors), equilibrated with 95% O_2_ at 5% CO_2_ and circulated at 2 ml/min at 34°C. A tungsten concentric bipolar stimulating electrode (FHC) was placed in the deep cortical layers, and the upper cortical layers were imaged with a 10x water-immersion objective (LUMPLANFL, Olympus) on an Olympus BX51 microscope. The stimulus pulses were generated through stimulus isolators ISO-Flex (A.M.P.I.) or S-88/SIU-5 (Grass Instruments). Stimulus intensity was set at 2x the resolvable threshold stimulation based on iGluSnFR imaging (Sup Fig 1.). Imaging was performed using a Zyla (Andor) camera (2048 X 2048 pixels, 4 X 4 spatial binning, 16-bit digitization, 10ms rolling shutter mode) and images were collected at 100 Hz temporal resolution, illuminated by a 470 nm LED (Thorlabs), using the endow-GFP filter cube (Chroma) and controlled by MicroManager (Edelstein et al. 2014).

**Figure 1.**
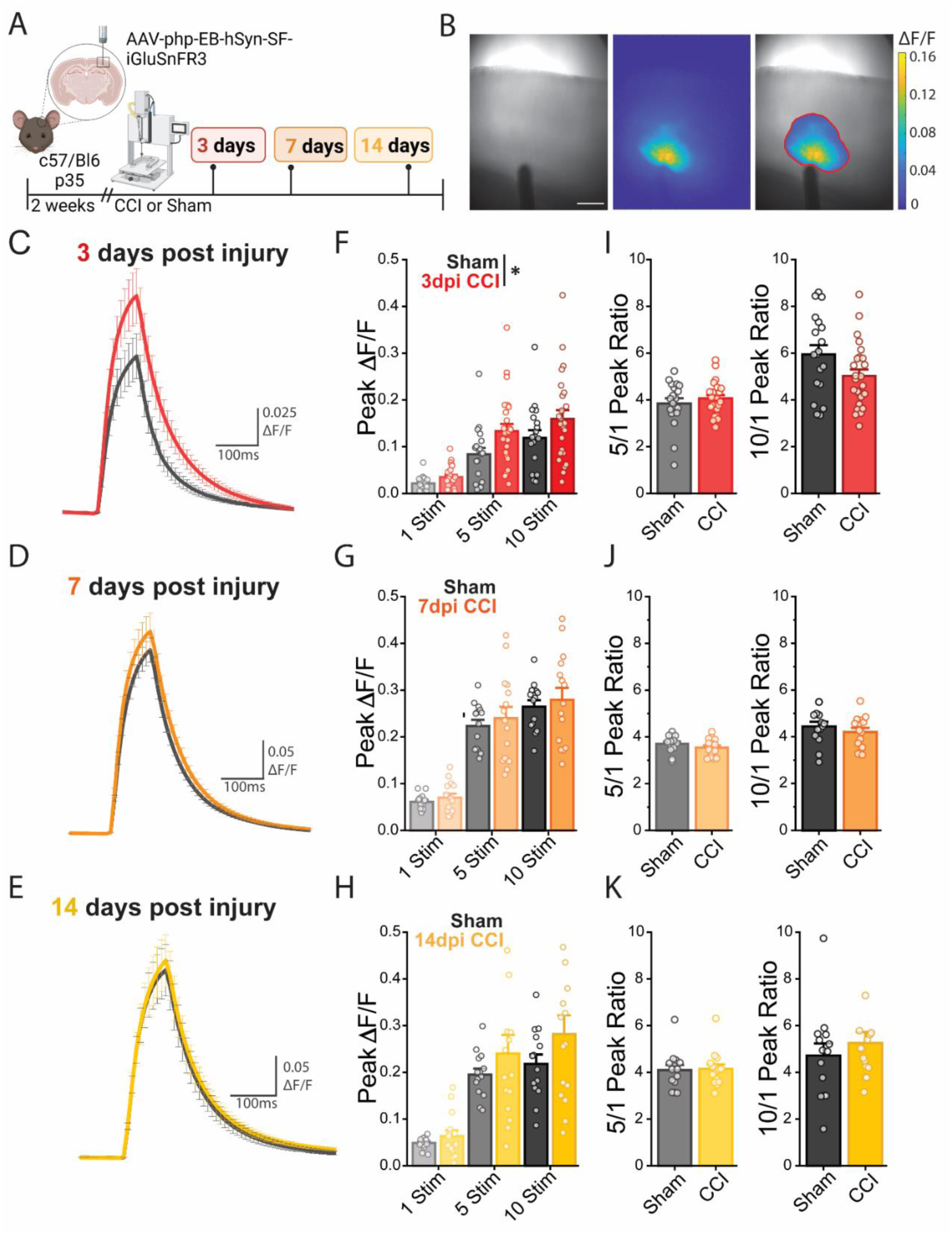
Stimulus evoked iGluSnFR signals are transiently elevated 3 days after CCI. **A.** Representative image of experimental workflow **B.** Representative 10X brightfield image showing placement of stimulus electrode in the deep cortical layers (*left*), evoked response iGluSnFR response (*center*), and the iGluSnFR ROI overlaid on the brightfield image (*right*). Scale bar, 250 μm. **C-E.** Average ΔF/F ten stimuli at 100Hz traces of iGluSnFR imaging comparing CCI to sham **C.** 3, **D.** 7, and **E.** 14 days after injury. **F.** Peak ΔF/F for all stimuli conditions 3 days after injury show increased peak glutamate 3 days after CCI for all stimuli conditions. LMM: p = 0.0476, *indicates P < 0.05 for differences between sham and CCI. Error bars indicate SEM. (Sham n= 4 mice, 19 ROIs; CCI n= 5 mice, 25 ROIs) **G, H.** Peak ΔF/F returns to sham **G.** 7 and (Sham n = 3 mice, 15 ROIs; CCI n = 3 mice, 15 ROIs) **H.** 14 days after CCI (Sham n = 3 mice, 15 ROIs; CCI n = 3 mice, 15 ROIs). **I-K.** Ratio of peak iGluSnFR response ratio to five to one and ten to one stimuli show no difference between sham and CCI for any time point.

### Drugs

Unless otherwise noted, all salts and glucose were obtained from Sigma-Aldrich. Drugs used in the study and their concentration were: APV (NMDA antagonist, 50μM, Tocris Bioscience) (Evans et al. 1982) and DNQX (AMPA antagonist, 20μM, Sigma) (Honore et al. 1988).

### Immunofluorescence

Brains were prepared for immunohistochemical analysis as previously described (Armbruster et al. 2016; Armbruster et al. 2022; Bottom-Tanzer et al. 2024). Briefly, mice we deeply anesthetized and transcardially perfused with ice cold PBS followed by 4% PFA. Brains we then post-fixed for 24 hours and then cryopreserved in 30% sucrose. Fixed brains were sectioned at 30 μm using a Thermo Fisher Microm HM 525 cryostat. Brain sections were blocked using blocking buffer (5% normal goat serum, 1% BSA, in PBS) for 1 h at room temperature. GLT-1 (1:1000, AGC-022-GP, Alomone Labs), GLAST (1:1000 AGC-021-Rb, Alomone Labs), Kir4.1 (1:400, APC-035, Alomone Labs), and green fluorescence protein (GFP, 1:1000; Abcam ab13970) antibodies were diluted in PBS with 2% Triton X-100 and 5% blocking buffer. Cortical sections were incubated with diluted primary antibodies overnight at 4°. Secondary antibodies (goat anti-guinea pig Cy3, goat-anti rabbit 488, Jackson ImmunoResearch Laboratories, goat-anti rabbit Cy3, Jackson ImmunoResearch) were diluted 1:500 in PBS with 5% blocking buffer and added to cortical sections for 2 h at room temperature. Slices were imaged with a Nikon A1R confocal microscope. Slices from 3 mice were immunolabeled for all experiments, with 2–4 slices per mouse visualized. For CCI-injured mice, ROIs were drawn on the cortex ipsilateral to injury beyond the lateral edge of the cavitary lesion superficial to CA3 unless otherwise detailed. ROIs were drawn in identical cortical regions in shams. All images were quantified using the mean fluorescence intensity within ROIs which were drawn and measured in ImageJ software (NIH).

### Imaging Analysis

#### Automated ROI detection

Custom MATLAB scripts were used for image processing and data analysis. A baseline fluorescence image was calculated by averaging 5 images occurring prior to stimulation onset. To identify regions of interest (ROIs), fluorescence change was calculated by subtracting the baseline fluorescence image from an average of 5 images during the peak iGluSnFR response. Pixels in this baseline-subtracted peak image that showed an iGluSnFR peak signal greater than 2X the standard deviation above the mean during the pre-stimulation time window were used to generate ROI masks. For each ROI mask, ΔF/F traces were generated by averaging across all pixels in the ROI at each time point. There was no significant difference of ROI size due to injury or day (Sup Fig 1.1). Baseline fluorescence (F) was quantified by averaging the raw imaging data prior to stimulation. To account for bleaching during imaging, every ΔF/F response was adjusted by subtracting the baseline and dividing by an imaging session in which no stimuli was delivered, as previously described (Armbruster et al. 2020; Armbruster et al. 2016). To determine iGluSnFR decays, ΔF/F traces were normalized by dividing all values by the peak response amplitude. Normalized mean ΔF/F traces were calculated by averaging across trials for each stimulation protocol and ROI. The post-stimulus decays were fit from the end of the stimulus to baseline with a mono-exponential decay. Peak values were calculated by finding the maximum ΔF/F_0_ value in the 20 images that followed stimulation.

#### Spatial analysis

In the spatial analysis, we established four distinct ROIs: one proximal to the injury (situated 20 μm from the injury site in layers III and IV), another distal to the injury, a third in the deep layer directly above the stimulus electrode, and a fourth in the superficial layers of the cortex. Importantly, these ROIs were non-overlapping, ensuring a clear distinction between them. Each ROI measured 50 μm by 50 μm and 20 μm were used to separate the proximal, deep, and distal ROIs. In this spatial data set we transformed our ratios using a log2 scale that transformed the data into a linear form, enabling the use of Linear Mixed Modeling (LMM) which is further detailed below.

#### Statistics

Statistical analyses were conducted using custom-written scripts in MATLAB and RStudio (2022.07.2 Build 576). Figures were created in OriginLabs and on BioRender.com. Normality was tested using the Shapiro-Wilk test. Linear mixed models (Aarts et al. 2014) were used to analyze the effects of injury (CCI vs. sham) and time after injury (3, 7, and 14 days) on glutamate dynamics, including peak responses and decay kinetics. Linear mixed modeling allows for the analysis of the effects of injury and stimulus number on peak responses and decay kinetics, while accounting for the fact that multiple measures come from the same animal and are thus not completely independent. We used the lme4 package in R to fit linear mixed models, incorporating fixed effects such as injury and their interactions. Including Mouse ID and slice number within that mouse as a random effect ensures the consideration of individual variability within the data. For peak analysis, peak glutamate response was the dependent variable. Fixed effects included injury, time after injury, and their interaction. Mouse ID was included as a random effect to account for repeated measures from the same animal. For decay kinetics, glutamate decay time constant was the dependent variable. Example code is as follows: > Glut3dpiDecaysLMM = lmer(Decay ∼ Stim + CCI + (1|Animal) + (1|Animal:Slice), data = Glut3dpiDecays, na.action = “na.omit”, REML = FALSE). For all models, statistical significance was defined as alpha < 0.05. When significant main effects or interactions were found, pairwise comparisons were made to determine differences between specific groups. Paired or two sample t-tests, nonparametric Wilcox-Signed rank test, or nonparametric Mann–Whitney test were used as appropriate. Samples sizes (slices/animals) are listed for each experiment in the figure labels or text, all experiments are from a minimum of 3 animals. Correction for multiple comparisons was tested using the Holm– Bonferroni method. Error bars indicate Standard Error of the Mean (SEM).

## RESULTS

### Stimulus-evoked glutamate responses are increased in amplitude transiently after CCI

We quantified extracellular glutamate dynamics after CCI using iGluSnFR imaging (Fig. 1A). Briefly, 2 weeks prior to injury, we injected (AAV.php.EB-hSyn-SF-iGluSnFR.A184V) (Marvin et al. 2018) into the brains of 5-week-old male C57BL/6 mice. We used a Leica Electromagnetic Impactor with a 3-mm-diameter piston to deliver a moderate CCI (3.5 m/s velocity, 400ms dwell time, and a 1 mm depth). This results in a focal contusion to the somatosensory cortex, without damage to underlying hippocampus. This protocol results in circuity hyperexcitability in acute cortical brain slices 2 weeks after injury (Graber and Prince 2004; Hunt et al. 2009; Koenig et al. 2019; Yang et al. 2010; Yang et al. 2021) and the development of post-traumatic epilepsy in a subset of animals (Bolkvadze and Pitkanen 2012; Guo et al. 2013; Hunt et al. 2009; 2010; Szu et al. 2020). Acute brain slices were prepared 3, 7, and 14 days following CCI, or sham injury, and iGluSnFR imaging was performed using a custom epifluorescence imaging setup (Fig. 1A). A tungsten concentric bipolar stimulation electrode was positioned in the deep cortical layers (Fig. 1B) to deliver a single stimulus or trains (5 or 10 stimuli at 100 Hz) of stimuli. In CCI tissue, the electrode was placed approximately 500 μm lateral to the site of injury in the white matter underlying cortex to activate ascending cortical axons. iGluSnFR signal was quantified within a region of interest (ROI) defined based on the iGluSnFR signal. Pixels that had a peak iGluSnFR signal 2 standard deviations greater than the mean iGluSnFR signal were included in the ROI (Sup Fig.1). To prevent post-synaptic glutamate receptor activation and circuit activity, brain slices were perfused with the AMPA receptor antagonist DNQX (20μm) (Evans et al. 1982) and the NMDA receptor antagonist APV (50μM) (Honore et al. 1988). This allowed quantification of glutamate signals arising solely from direct electrical stimulation, with no contribution of subsequent evoked or spontaneous circuit-level activity.

We found that the peak stimulus-evoked iGluSnFR signal was significantly increased 3 days after CCI from 0.0822 ± 0.012 to 0.122 ± 0.014 after 10 stimulations (Fig. 1C, F). This change was transient and when cortical brain slices from mice 7 (Fig. 1D, G) and 14-days (Fig. 1E, H) after injury were examined, there was no difference between CCI and sham-injured animals in peak iGluSnFR response. Subsequently, we explored how the number of stimuli affected the amount of glutamate released, as assayed by iGluSnFR peak ratios. We calculated the ratio of iGluSnFR peaks between 1 and 5 stimuli and 1 and 10 stimuli to assay the linearity of stimulus-dependent glutamate release. For both sham and CCI injured animals, there were sub-linear increases in iGluSnFR signal with increasing stimuli, but there was no difference between sham and CCI at any time point (Fig. 1I-K). Together this shows that CCI transiently increases peak stimulus-evoked iGluSnFR signal without affecting the input-output relationships.

### Glutamate uptake is slowed 3 days after CCI, but activity-dependent inhibition of glutamate uptake is unchanged

To test whether glutamate uptake is compromised after CCI, we analyzed the decay times of stimulus-evoked iGluSnFR signals. We have previously shown that changes in iGluSnFR decay times reflect alterations in astrocyte glutamate uptake (Armbruster et al. 2016; Armbruster et al. 2022). iGluSnFR decay times were calculated by fitting the post-stimulus recovery with an exponential decay and compared across stimuli and conditions. We found that CCI significantly slowed iGluSnFR decay times 3 days after injury to 86.7 ms ± 6.18 from 60.18 ms ± 4.07 for sham animals (10 stimuli; Fig. 2A, D). This slowing of glutamate uptake was seen for all stimuli numbers examined 3 days after CCI. Similar to changes in stimulus-evoked iGluSnFR amplitude, changes in decay time were also transient. At 7 (Fig. 2B, E) and 14 days (Fig. 2C, F) after CCI, iGluSnFR decay times were similar in sham and CCI mice. We then quantified whether CCI altered activity-dependent inhibition of glutamate uptake. We recently reported that neuronal activity rapidly modulates glutamate uptake (Armbruster et al. 2016) via voltage-dependent inhibition of EAATs (Armbruster et al. 2022) mediated by neuronal release and astrocyte uptake of K^+^. We calculated the ratio of iGluSnFR decay times between 1 and 5 stimuli and 1 and 10 stimuli to assay the magnitude of activity-dependent inhibition of glutamate uptake. In both sham and CCI animals, glutamate uptake was reduced with trains of stimuli (Fig. 2G-I), consistent with previous reports (Armbruster et al. 2016; Armbruster et al. 2022; Barnes et al. 2020). There were no differences, however, in the magnitude of stimulus-dependent slowing between CCI and sham groups at any timepoint. This shows that there was a significant decrease in glutamate uptake 3 days after injury but no changes in activity-dependent inhibition of glutamate uptake.

**Figure 2.**
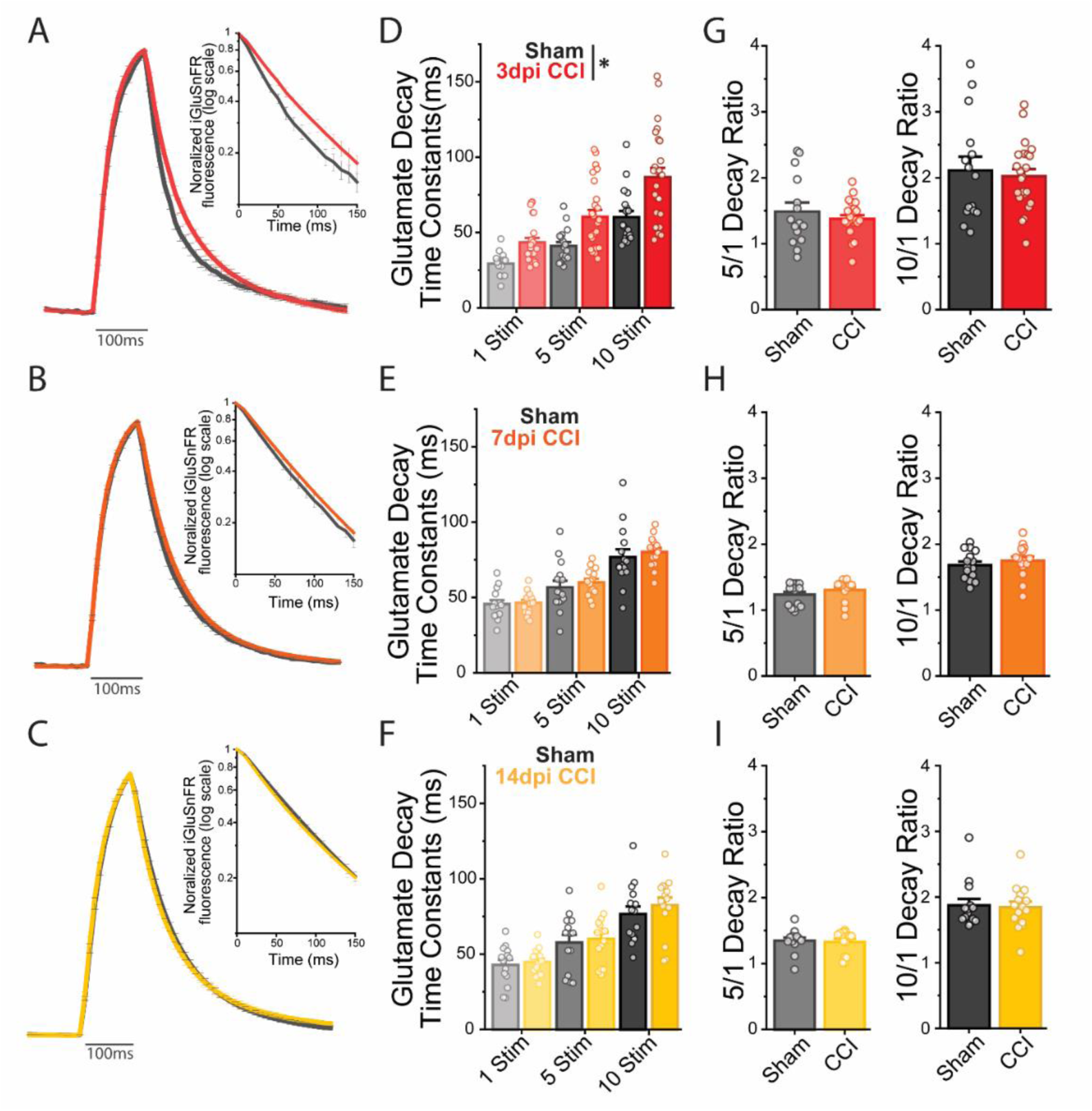
Glutamate uptake is slowed 3 days after CCI, but activity-dependent inhibition of glutamate uptake is unchanged. **A-C.** Normalized iGluSnFR responses to 10 stimuli delivered at 100 Hz show slowed glutamate uptake. **A.** 3 days after CCI compared to sham. **B.** By 7 and **C.**14 days, normalized iGluSnFR responses have similar decay times between CCI and sham animals. **D-F.** Quantified iGluSnFR decay times show increased glutamate decay time constants **D.** 3 days after CCI. LMM: p = 0.03557, *indicates P < 0.05 for differences between sham and CCI. Error bars indicate SEM. (Sham n= 4 mice, 19 ROIs; CCI n= 5 mice, 25 ROIs) **E.** 7 and (Sham n = 3 mice, 15 ROIs; CCI n = 3 mice, 15 ROIs) **F.** 14 days after CCI (Sham n = 3 mice, 15 ROIs; CCI n = 3 mice, 15 ROIs) Decay times return to sham **G-I.** Activity dependent inhibition of glutamate uptake is not changed after CCI for anytime point.

### CCI does not significantly alter iGluSnFR expression

To test whether CCI altered expression of iGluSnFR, we performed CCI and sham surgeries on animals focally injected with iGluSnFR. While this is the first study to utilize iGluSnFR in a TBI model, other studies using GCaMP in the CCI model show no significant difference in expression following injury (Bottom-Tanzer et al. 2024). We used immunohistochemical labeling of GFP and epifluorescence imaging to quantify iGluSnFR expression. We examined 3 days post-surgery, as this is the time most likely to show changes in expression and is the time in which we report changes in glutamate uptake kinetics. Brains were fixed and prepared for immunohistochemical analysis of the reporter expression using an anti-GFP antibody. We found no significant difference in GFP between CCI and sham mice (Fig. 3C). This supports that any changes in iGluSnFR activity in the imaged areas are not due to differences in reporter expression, but rather due to changes in glutamate signaling.

**Figure 3.**
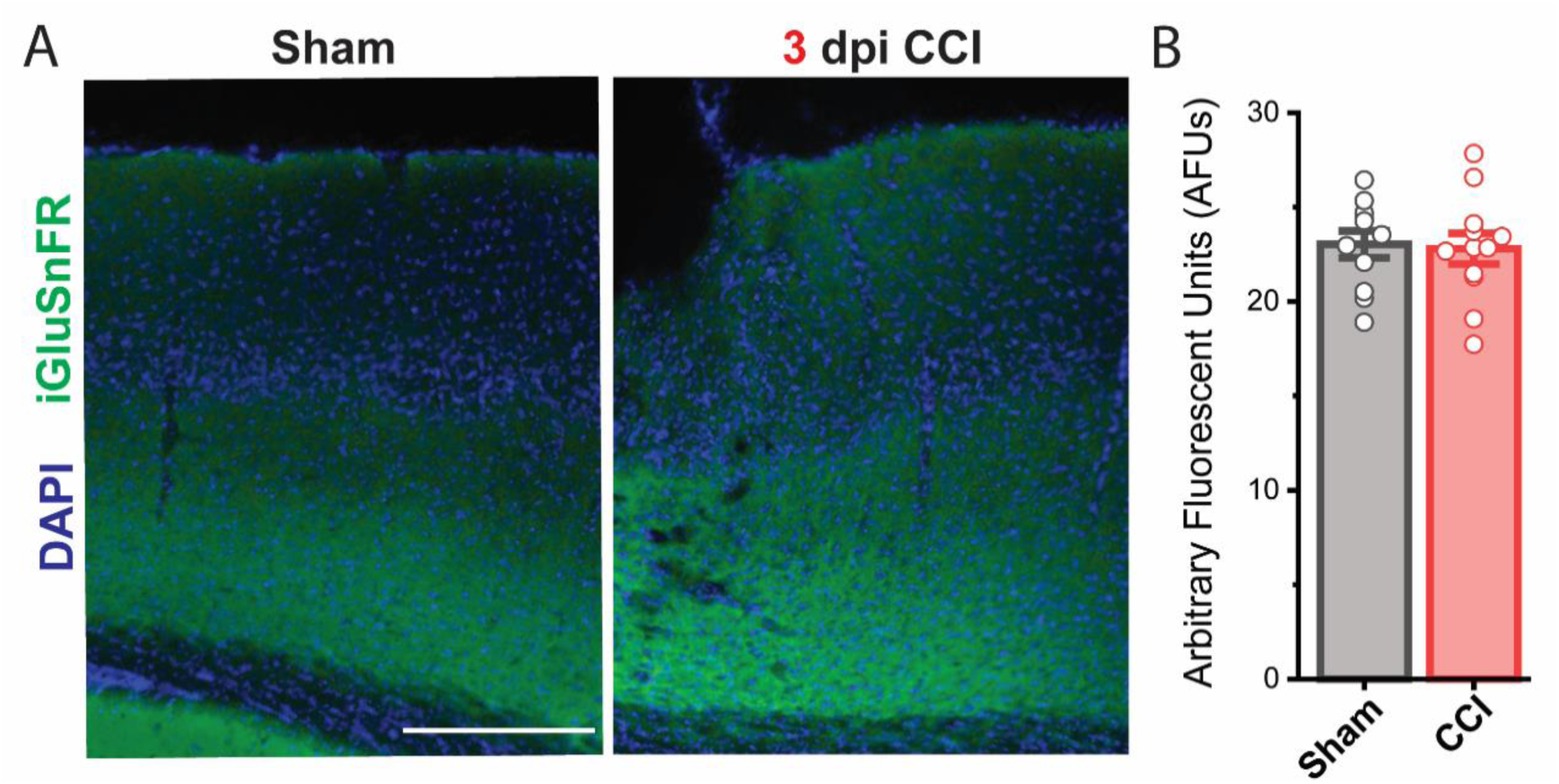
iGluSnFR expression is not significantly altered by CCI. **A.** Representative fluorescent images of sham and CCI injured mice 3 days post-surgery immunolabeled for GFP (green). Scale bar, 100μm. The CCI representative image is perilesional. **B.** Mean GFP fluorescence was not significantly affected by CCI. Error bars indicate SEM: 3 days after injury (Sham n= 4 mice, 3 slices per mouse; CCI n= 4 mice, 3 slices per mouse).

### GLT-1, but not Kir4.1 or GLAST, is transiently reduced after CCI

Because CCI increased iGluSnFR decay times and peaks, but not stimulus-dependent slowing of glutamate uptake, we quantified immunolabeling of GLT-1, GLAST, and Kir4.1 following CCI to determine if functional changes are associated with changes in expression. Based on our previous findings (Armbruster et al. 2016; Armbruster et al. 2022), we hypothesized that basal glutamate uptake is established by EAAT expression levels, while activity-dependent slowing of glutamate uptake is driven by Kir4.1 expression. Because CCI prolonged iGluSnFR decay times 3 days after injury but had no effect on activity-dependent slowing, we suspected that EAAT levels would be decreased, while Kir4.1 levels would be unaltered (Armbruster et al. 2016; Armbruster et al. 2022). Immunoreactivity for each protein was quantified by measuring average intensity within the peri-injury area of the somatosensory cortex (Fig. 4A), corresponding to the area of cortex that was imaged in during iGluSnFR imaging. Consistent with our hypothesis, GLT-1 immunoreactivity was significantly reduced 3 days after CCI but returned to sham levels by 7 days post-injury (Fig. 4B, D). Kir4.1 immunoreactivity was unaltered in sham and CCI mice at all timepoints (Fig. 4C, E).

**Figure 4.**
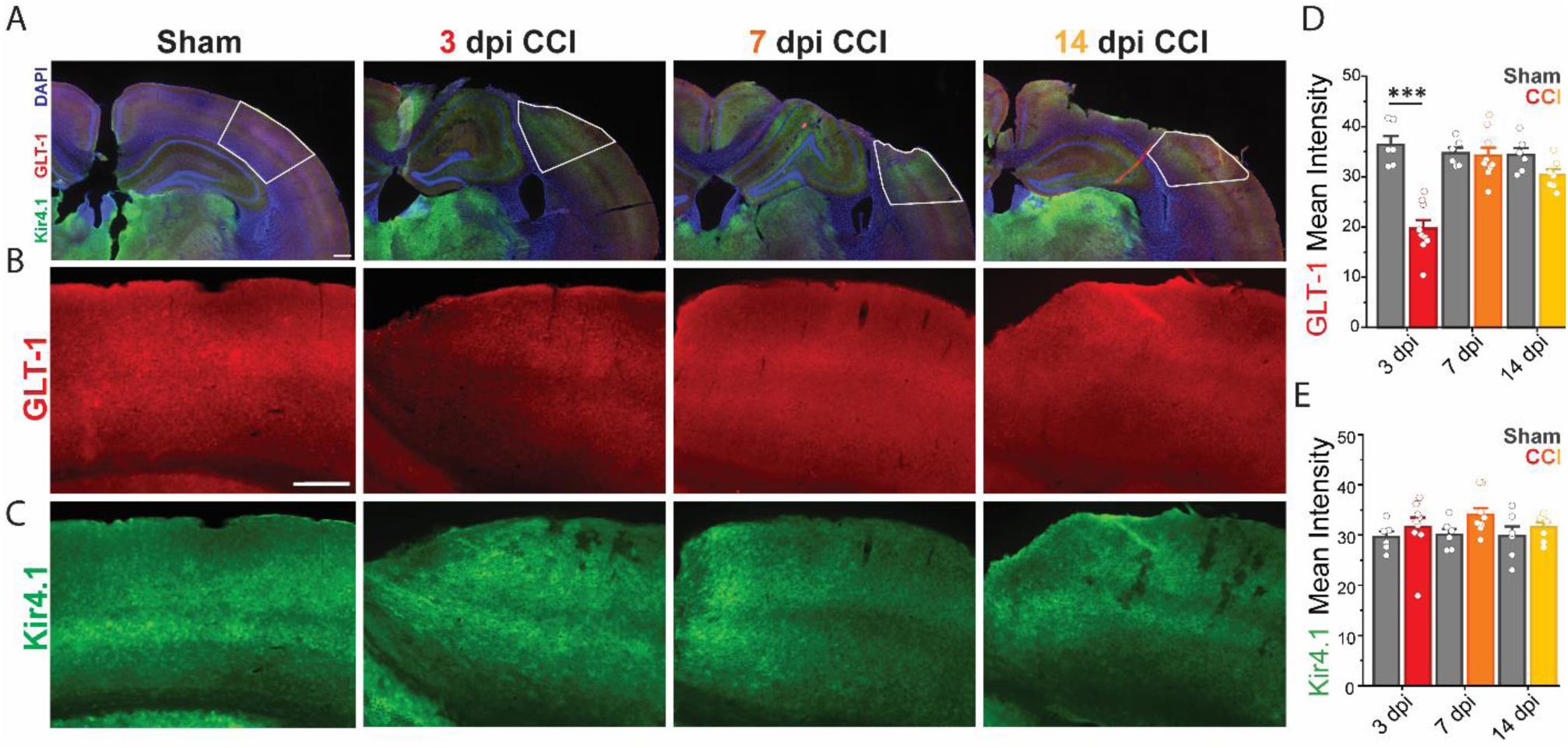
GLT-1, but not Kir4.1, levels are transiently reduced after CCI. **A.** Representative images of sham and CCI highlighting analyzed ROIs (white boxed area). Scale bar = 200μm **B.** GLT-1 and **C.** Kir4.1 immunolabels for sham and CCI; scale bar = 100μm. **D.** Mean intensity analysis shows a significant reduction of GLT-1 3 days after CCI. **E.** Mean intensity analysis shows no change in Kir4.1 after injury for any time point. One-way repeated-measures ANOVA and Tukey’s posttest: sham versus CCI 3 days after injury, p = 1.9x10^-5^; ***indicates P < 0.001 (all conditions n = 3 mice, 9 slices).

After observing a dramatic loss of GLT-1 3 days after injury, we next quantified if GLAST, another astrocyte EAAT. GLAST is highly expressed during development (Kugler and Schleyer 2004) and contributes to glutamate uptake in both the adult and developing brain (Chaudhry et al. 1995; Peghini et al. 1997). Surprisingly, we found no significant change in GLAST levels in CCI animals, compared to shams, 3 days after injury (Fig 5A-C) even though GLT-1 was altered by injury. These results show specific downregulation of GLT-1 as a mechanism of glutamate dysfunction after TBI, with no change in GLAST or Kir4.1 levels and supports our model for how EAATs and Kir4.1 work to shape the dynamic nature of glutamate uptake.

**Figure 5.**
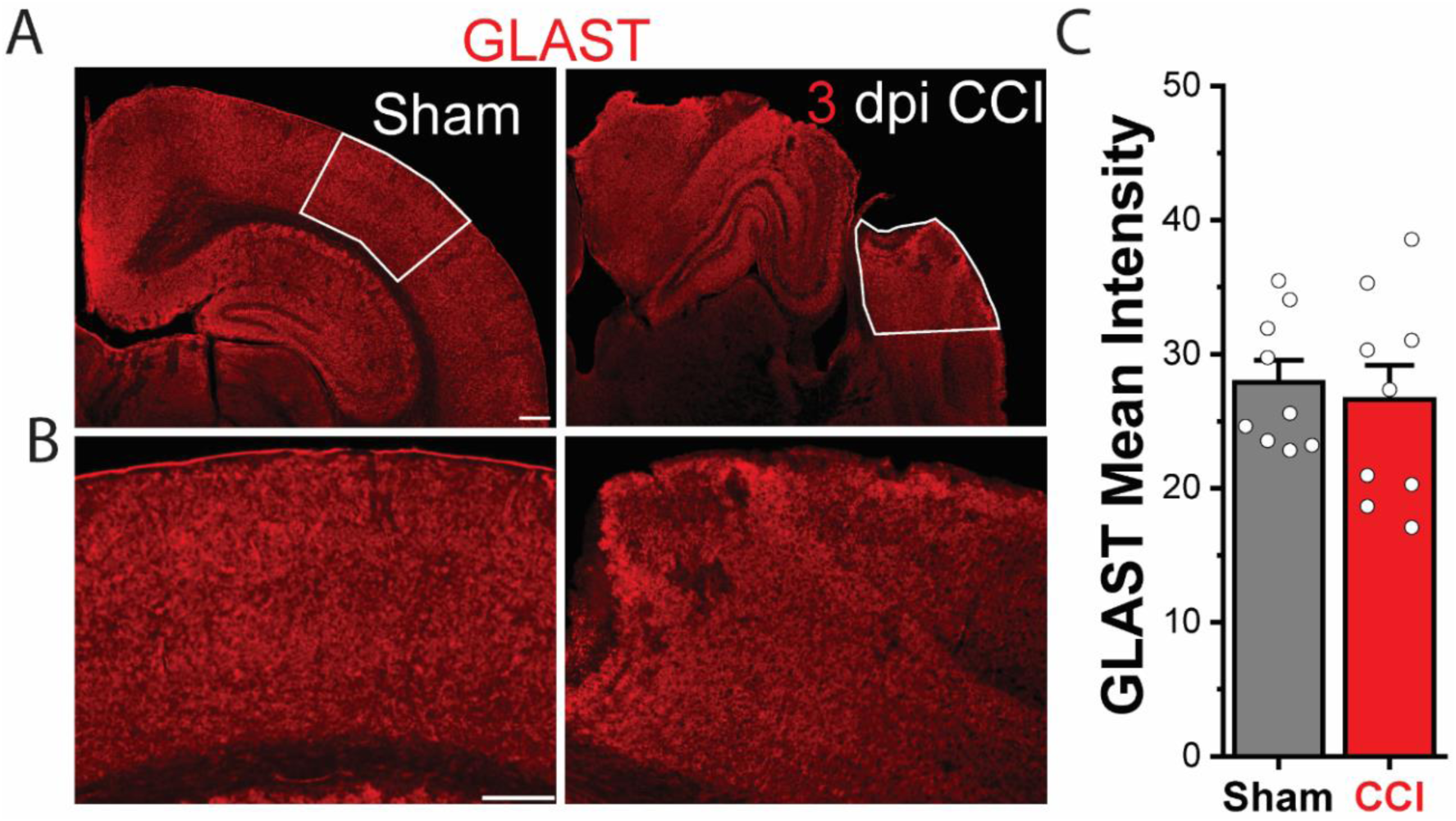
GLAST levels are unchanged 3 days after CCI. **A.** Representative images of sham and CCI immunolabeled with an anti-GLAST antibody 3 days after CCI and highlighting analyzed ROIs (white boxed areas). Scale bar = 200μm. **B.** Representative confocal IHC images immunolabeled for GLAST for sham and CCI 3 days after injury: scale bar = 100μm. **C.** Mean Intensity of GLAST for the ROIs outlined in **A.** 3 days after injury shows no significant changes in GLAST immunolabeling (all conditions: n = 3 mice, 9 slices).

### CCI induces spatially heterogenous changes in peak iGluSnFR amplitude, but glutamate uptake is spatially uniform

TBI causes spatially and temporally heterogenous disruptions to cortical circuits (Budinich et al. 2013; Song et al. 2016; Zhao et al. 2015). We were curious whether the changes in extracellular glutamate dynamics that we report are spatially-uniform or heterogenous. Changes in glutamate dynamics proximal to the site of injury, versus distally, would be particularly relevant to peri-injury specific pathology. In addition, changes in layer-specific neuronal projections have been seen in brain injury (Cantu et al. 2015; Frankowski et al. 2022) and could impact circuit function. To quantify regional differences in glutamate release and uptake dynamics following CCI, we defined and analyzed four distinct regions of interest (ROIs) in the injured and sham cortex. Within the stimulated area of cortex (10 stimuli delivered at 100 Hz), we examined the laminar distribution of iGluSnFR signal (deep cortical layers IV-VI vs. superficial cortical layers I-III). We also examined iGluSnFR signal proximal the CCI injury in the deep cortical layers (medial to the stimulation electrode, <100 μm from the injury in layers III-IV) and distal to the injury (lateral to the stimulation electrode, ∼500 μm from the injury in layers III-IV) (Fig. 6B, C). The size of each ROI was 250 μm X 250 μm and neighboring ROIs were separated by 20 μm to ensure no spatial overlap. Peak iGluSnFR response was increased in superficial layers (relative to deep layers) (Fig. 6D) and adjacent to the site of injury (relative to distal to the site of injury) 3 days after CCI, as compared to shams (Fig. 6E). Interestingly, changes in glutamate uptake, as assayed by iGluSnFR decay times, were similar in sham and CCI as well as all regions, (Fig. 6F,G). These results demonstrate that CCI causes spatially unique changes in peak glutamate response, but that changes in glutamate uptake are spatially uniform 3 days after injury (Fig. 6D-G).

**Figure 6.**
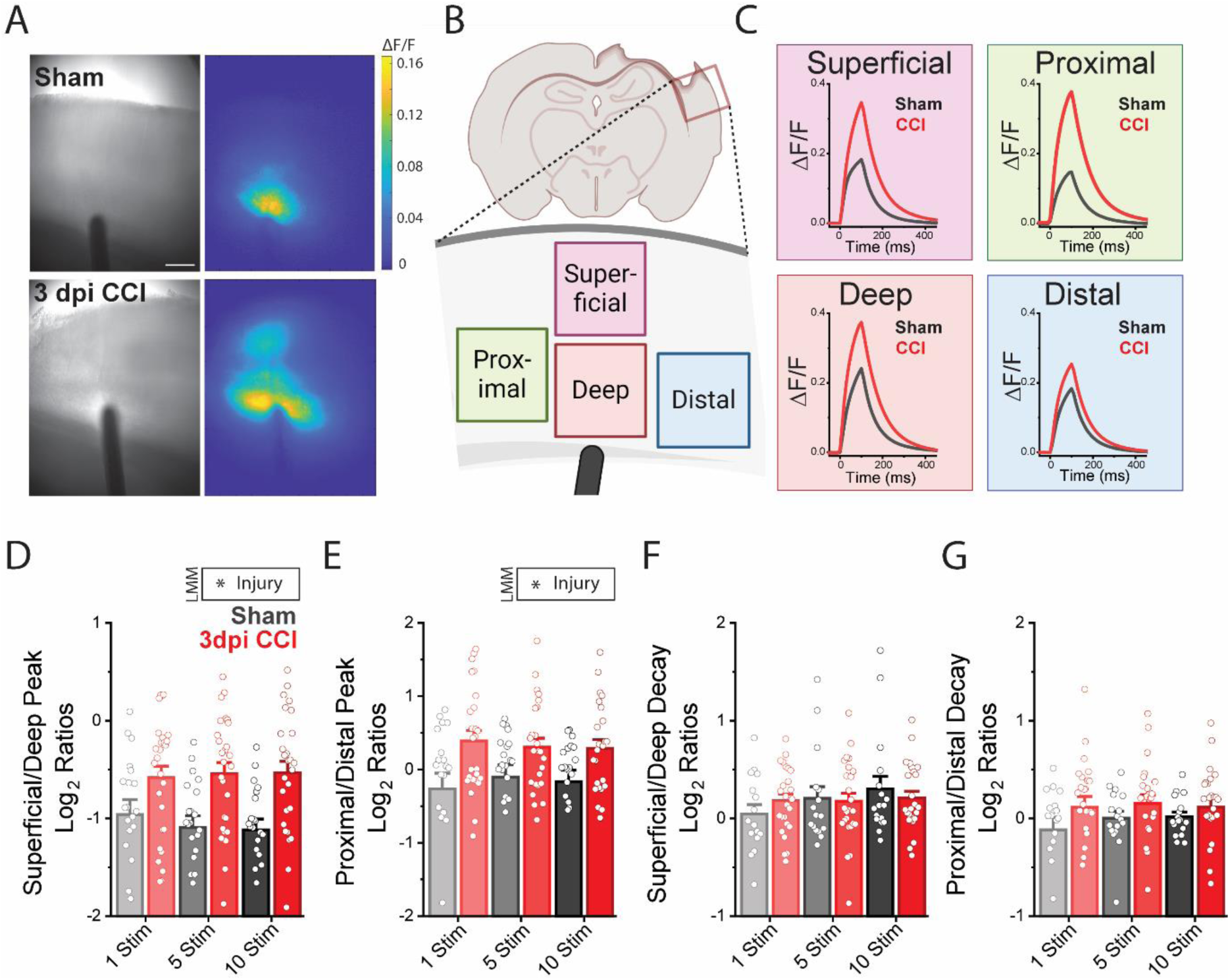
CCI-induced changes in peak iGluSnFR signal, but not glutamate uptake, are spatially heterogenous. **A.** Brightfield and peak response images for sham and CCI 3 days after surgeries with corresponding iGluSnFR images. Scale bar, 250 μm. **B.** Schematic showing the placement of ROIs for spatial analysis. **C.** Ten stimuli iGluSnFR responses in each ROIs show increased peak glutamate for all ROIs in comparison to sham. **D.** The superficial over deep peak ratios demonstrate increased peak glutamate compared to sham 3 days after injury for all stimuli conditions. LMM: p = 0.01675, *indicates P < 0.05 for differences in injury. Error bars indicate SEM: 3 days after injury (Sham n= 4 mice, 19 ROIs; CCI n= 5 mice, 25 ROIs). **E.** Similarly, the proximal over distal peak ratios demonstrate increased peak glutamate. LMM: p = 0.0397 *indicates P < 0.05 for differences in injury. **F.** Superficial over deep decays show no difference due to injury. **G.** No significant difference is seen between CCI and sham animals when comparing Proximal/Distal Decay ratios.

### Injury causes homogeneous loss of GLT-1 three days after injury

Because iGluSnFR analysis suggested spatially-uniform changes in glutamate uptake after injury (Fig. 6D), we next asked if changes in the expression of GLT-1 were also spatially uniform. Using regional ROIs analysis, similar to our regional iGluSnFR analysis (Fig. 7A), we quantified GLT-1 immunolabeling throughout the CCI- and sham-injured cortex using 250 μm X 250 μm ROIs separated by 20 μm. We found a significant reduction in GLT-1 for 3 days after injury (Fig. 7B-D) but did not observe any significant changes in Kir4.1 expression following injury in any region or at any timepoint (Fig. 7C-D). This supports our iGluSnFR imaging analysis of glutamate uptake and shows that while there are many spatially-distinct changes after TBI, changes in GLT-1 are spatially-uniform within the area analyzed. Together this suggests that spatially heterogenous changes we report in peak iGluSnFR response caused by CCI are likely not driven by changes in GLT-1, but rather likely arise from changes in glutamate release.

**Figure 7.**
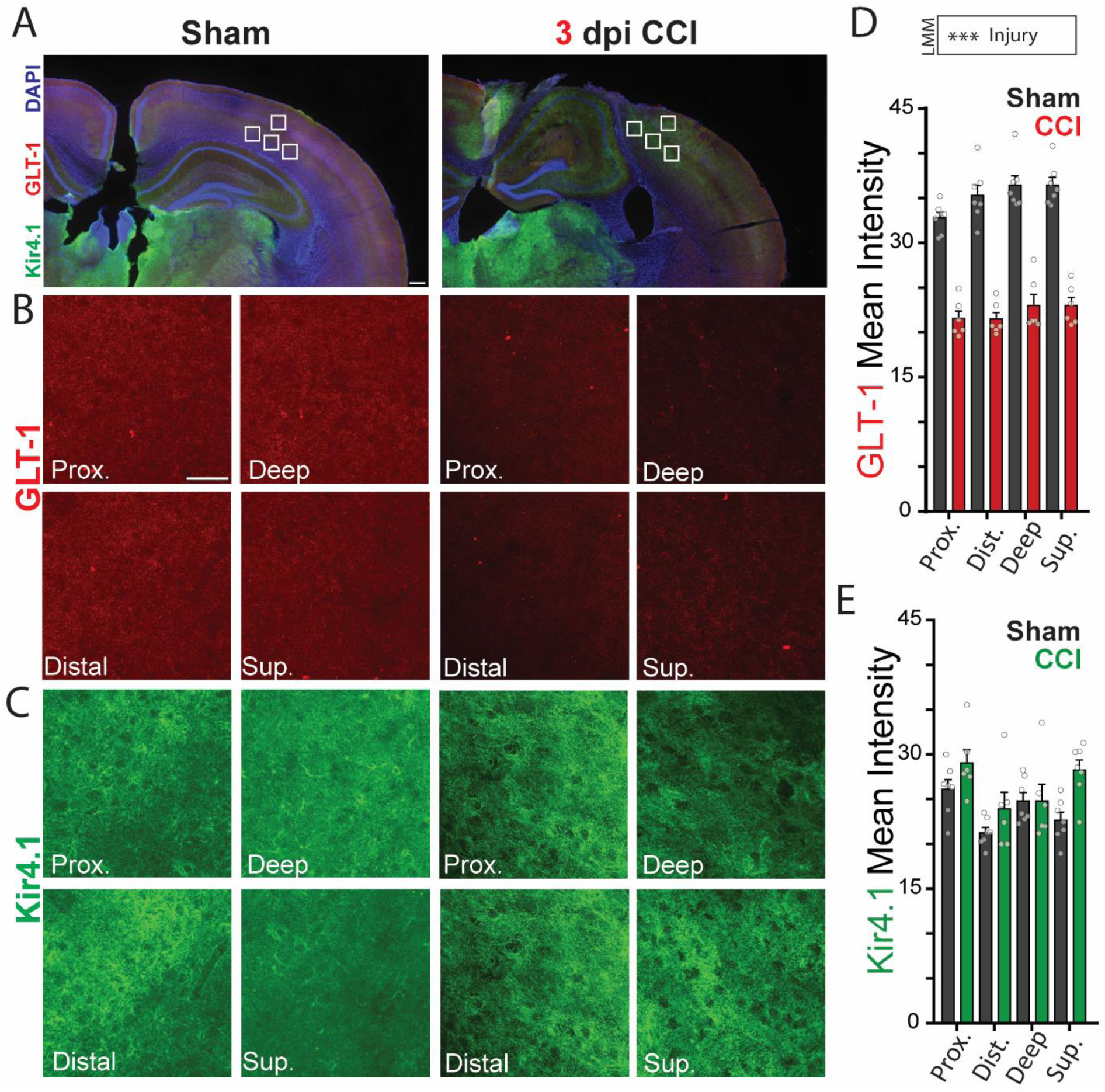
Injury causes homogeneous loss of GLT-1 three days after injury. **A.** Representative images of sham and CCI highlighting analysis ROIs (white boxed area). Scale bar = 200μm **B.** Immunolabeling of GLT-1 in sham and CCI for each ROI; scale bar = 20μm. **C.** Immunolabeling of Kir4.1 in sham and CCI for all ROIs. **D.** Mean intensity of GLT-1 immunolabeling show a significant reduction of GLT-1 3 days post injury. LMM: Injury p=1.96e-05 for differences between sham and CCI; *** indicates P <0.0001. Error bars indicate SEM. (all conditions: n= 3 mice, 9 slices) **E.** Mean Intensity of Kir4.1 by ROI and day shows no significant changes in Kir4.1 (all conditions: n= 3 mice; 9 slices).

## DISCUSSION

Our study used a combination of iGluSnFR imaging and immunohistochemical analysis to examine how changes in astrocytic EAATs and Kir4.1 contribute to injury-induced disruption of extracellular glutamate dynamics. We found that CCI transiently enhanced glutamate signaling, increasing iGluSnFR peak signal and prolonging decay times 3 days post-injury. Further, we found corresponding decreases in GLT-1 expression 3 days post-injury. These changes were transient and by 7 days, iGluSnFR responses and GLT-1 immunolabeling were similar in CCI and shams (Fig 8.).

**Figure 8.**
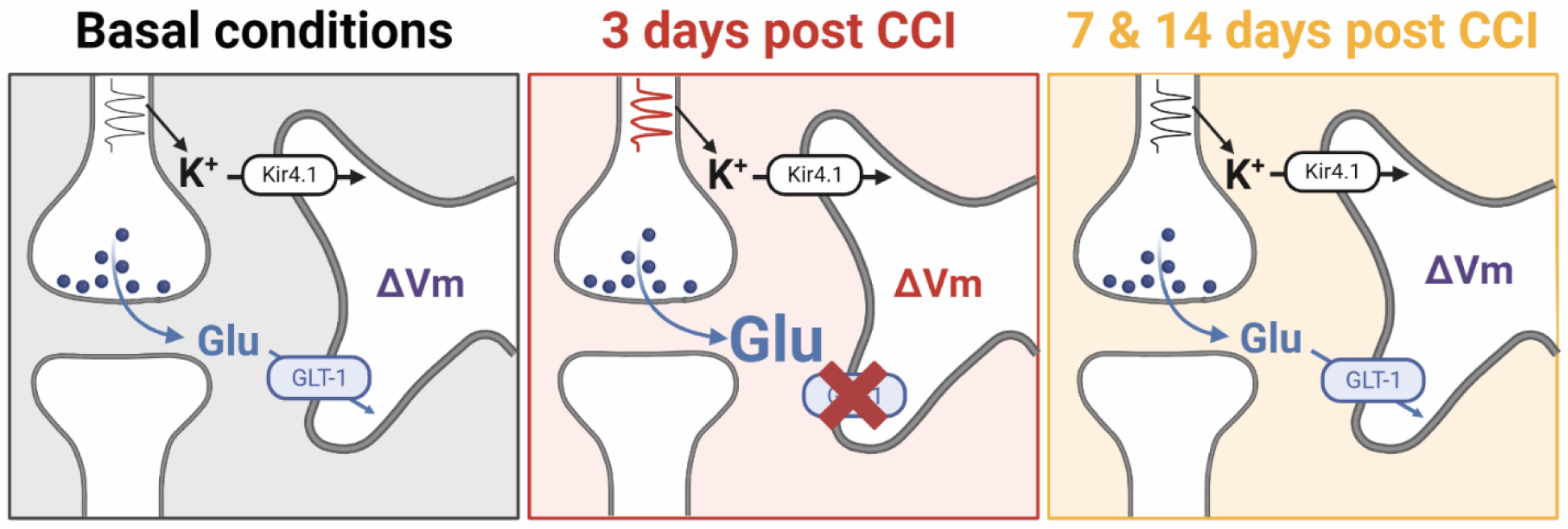
TBI transiently alters GLT-1 expression and glutamate uptake. During normal conditions, K^+^ and glutamate are released from the presynaptic neuron. Following CCI, astrocytic GLT-1 expression is decreased, leading to increased glutamate in the extracellular space. By 7 and 14 -days after injury, GLT-1 levels return, and glutamate uptake is restored.

Based on our previous study examining activity-dependent inhibition of glutamate uptake (Armbruster et al. 2016; Armbruster et al. 2022), we suspected injury-induced decreases in Kir4.1 expression would exaggerate activity-induced suppression of glutamate uptake. In other studies, Kir4.1 expression (Gupta and Prasad 2013; Shandra et al. 2019; Zurolo et al. 2012) is decreased after injury, which we predicted would enhance activity-induced suppression of glutamate uptake. In our experiments, however, we did not see loss of Kir4.1 after CCI and activity-induced inhibition of glutamate uptake was not altered by TBI. This result, while different from other studies showing injury induced decreases in Kir4.1 (Gupta and Prasad 2013; Shandra et al. 2019), supports our model in which Kir4.1 and GLT-1 interact to dynamically shape glutamate uptake (Fig. 8).

Using iGluSnFR imaging allowed us to also examine how injury affects glutamate dynamics across the cortex. While the peak iGluSnFR signal was spatially-heterogeneous after CCI, with increases seen in superficial cortical layers and proximal to the site of injury, the rate of glutamate uptake was spatially-uniform. This contrasts with our previous observations, where we noted increased glutamate in deep cortical layers proximal to the injury, but no increase in superficial layers. This discrepancy could largely be due to variations in imaging techniques and our use of post-synaptic NMDA and AMPA inhibitors to isolate astrocyte glutamate uptake in the current study vs circuit changes in our previous work (Cantu et al. 2015). Consistent with our glutamate uptake findings, GLT-1 immunolabeling was spatially-uniform within all regions examined. Together this data suggests that regional heterogeneity in peak iGluSnFR response is not due to focal loss of astrocyte glutamate uptake, but rather likely arises due to altered glutamate release, presynaptic function (Reeves et al. 2000), and/or axonal reorganization (Jenkins and Merzenich 1987). TBI is known to alter axonal structure (Johnson et al. 2013), glutamate synaptic function (Thomas et al. 2012), and axonal excitability (McNamara et al. 2010; Thomas et al. 2012) consistent with potential changes in glutamatergic output.

Our immunolabeling experiments show a significant reduction in GLT-1 after TBI, suggesting a significant disruption of glutamate homeostasis (Danbolt 2001) which could promote pathological excitatory synaptic transmission and circuit activity following TBI. This is consistent with multiple studies that report reduced GLT-1 expression and function after TBI (Goodrich et al. 2013; Gupta and Prasad 2013; Hinzman et al. 2012; Rao et al. 1998; Shandra et al. 2019; van Landeghem et al. 2001; Yi et al. 2005). These changes, however, vary across TBI models, time after injury, and region examined. CCI studies report early loss of GLT-1 protein in the ipsilateral cortex starting 4-6 hours after injury and persisting past 72 hours (Rao et al. 1998). Previously, we demonstrated using western blot that 14 days after CCI, GLT-1 and GLAST returned to sham levels(Cantu et al. 2015). Studies utilizing the lateral fluid percussion model show no change in GLT-1 in the first 24 hours but subsequent decreases when examined 7 days post injury (Goodrich et al. 2013; Yi et al. 2005). The impact acceleration (weight drop) injury model shows a decrease in both GLT-1 and Kir4.1 14 days after injury in the hippocampus and cortex (Shandra et al. 2019). Studies from human postmortem tissue find a decrease in astrocytic glutamate transporter expression following TBI (Ikematsu et al. 2002; van Landeghem et al. 2006), with a dramatic loss of GLT-1 that begins 24 hours after injury and persists for approximately 7 days. At later survival times, GLT-1 returns to normal levels.

Several potential mechanisms have been linked to injury-associated changes in GLT-1. Studies in cultured cortical astrocytes have shown that exposure to high levels of extracellular glutamate (20mM) decreased GLT-1 expression levels by 25% and 40% following 24- or 72 hour exposure times, respectively (Lehmann et al. 2009). Although the mechanisms linking glutamate and decreased GLT-1 levels are unknown, this suggests that the decrease in GLT-1 may occur due to increased extracellular glutamate after TBI (Chamoun et al. 2010; Katayama et al. 1990; Sowers et al. 2021). Conversely, during development metabotropic glutamate receptor activation promotes GLT-1 (Morel et al. 2014) expression. Other potential mechanisms include alterations in microRNAs (miRNAs) known to modulate GLT-1 (Meng et al. 2017; Pan et al. 2017), caspase-3 cleavage of GLT-1, and PCK activation (Kalandadze et al. 2002), all of which have all been reported following injury. It is known that miRNAs are critical regulators of gene expression and have been shown to regulate GLT-1 expression levels in an epilepsy model (Wang et al. 2003). Specifically, miR-181c-5p was shown to be released in neuronal extracellular vesicles and can worsen epileptic seizures by inhibiting the PKCδ/GLT-1 signaling axis (Kalandadze et al. 2002), which led to a decrease in glutamate uptake by astrocytes. Caspase-3 has been shown to cleave and inactivate GLT-1, impairing its function (Boston-Howes et al. 2006; Gibb et al. 2007; Rosenblum et al. 2017). Multiple studies have demonstrated elevated caspase-3 following traumatic brain injury (Beer et al. 2000; Chen et al. 2004b), suggesting another potential mechanism for a reduction in GLT-1 levels.

TBI also initiates inflammatory responses (Lagraoui et al. 2012; Shohami et al. 1997) which could impair GLT-1 expression and function. Multiple studies have demonstrated that both IL-1β (Mandolesi et al. 2013) and TNF-α (Carmen et al. 2009; Tolosa et al. 2011) signaling reduce EAAT expression (Mandolesi et al. (2013); (Takaki et al. 2012). Cerebral spinal fluid from TBI patients show increased concentrations of both IL-1β and TNF-α (Hayakata et al. 2004) 24 hours after injury, a period in which GLT-1 is reduced in TBI patients. Levels gradually decrease overtime, thereafter, suggesting a potential neuroinflammatory mechanism mediating loss of GLT-1. Therapeutic interventions aimed at restoring GLT-1 expression and function could hold promise in mitigating the neurophysiological consequences of TBI. For example, ceftriaxone enhances GLT-1 expression and has been used in the injured brain to improve glutamate uptake and functional recovery (Goodrich et al. 2013).

Interestingly, even though we see a dramatic reduction in GLT-1 expression, we observed a relatively small increase in glutamate decay times. This could be due to the fact that the iGluSnFR signal is a combination of the glutamate signal itself and the kinetics of iGluSnFR. Glutamate clearance is very fast (1-10 ms), whereas the iGluSnFR sensor is slower than the glutamate clearance itself. Since our imaging data convolves clearance and the sensor response, we potentially underestimate changes in glutamate clearance (Armbruster et al. 2020). Additionally, CCI reduces GLT-1 immunolabeling, but GLAST levels are unchanged. Because of this differential regulation of these two important astrocyte EAATs after CCI, remaining GLAST may be able to compensate for lost GLT-1. Our previous studies show that pharmacological block of GLT-1 alone causes relatively small changes in iGluSnFR decay kinetics, while blocking all EAATs has a much larger effect (Armbruster et al. 2016). In addition, *in vitro* studies show increased GLAST functional activity without an increase in protein expression (Susarla et al. 2004) suggesting potential enhanced GLAST function when GLT-1 is lost. Interestingly, previous studies have shown increased (Gilley and Kernie 2011), decreased (Rao et al. 1998), and no change (Yi and Hazell 2006) in GLAST levels following TBI. Similarly, to Kir4.1 and GLT-1, these differences are likely due to difference in time after injury, region examined, and model of TBI. Several signals govern the regulation of both GLT-1 and GLAST, such as NF-κB (Karki et al. 2015) and β-catenin (Lutgen et al. 2016). However, subtype specific regulation of EAATs by N-myc (Gupta and Prasad 2014) could explain the differential regulation of GLT-1 versus GLAST following TBI. Exploring these mechanisms could offer new perspectives on how TBI selectively effects GLT-1 versus GLAST and how those signaling pathways could be harnessed to treat injury.

While our study provides valuable insights into the spatiotemporal dynamics of glutamate after CCI, there are caveats that must be noted. Our study exclusively focused on the perilesional cortex ipsilateral to injury, so changes that occur distally or contralaterally to the injury site could also occur. Additionally, we only examined 3, 7, and 14 days post-TBI. Given that TBI evokes dynamic molecular and cellular responses, examining earlier time points could yield additional insights into the dynamics of glutamate transporter alterations post-injury. Moreover, the accelerated pace of physiological changes in mouse models may not accurately mirror the more protracted temporal patterns observed in human TBI cases. Furthermore, since CCI did not alter Kir4.1 levels, and we hypothesized that Kir4.1 modulates the activity-dependent slowing of glutamate uptake (Armbruster et al. 2016), we may be able to further probe this hypothesis using other TBI models that are known to evoke changes in Kir4.1. Finally, our study focused solely on male mice; whether the changes we report here also occur in females remains to be seen. Current research suggests that there are sex differences in the response to TBI with variations in outcomes and mechanisms between male and female subjects (Doran et al. 2019; Jullienne et al. 2018; Kupina et al. 2003; Villapol et al. 2017). Exploring these differences further could enhance our understanding of glutamate dynamics post-TBI and inform more tailored treatment approaches.

Those caveats aside, our study shows dynamic changes in glutamate neurotransmission after injury, demonstrates that injury does not affect either Kir4.1 or activity-dependent inhibition of glutamate uptake, and causes spatially-specific changes in peak glutamate responses but spatially-uniform changes in glutamate uptake. Furthermore, our study employed iGluSnFR as a novel tool to investigate glutamate dynamics following CCI. By utilizing this cutting-edge imaging technique, we were able to capture real-time changes in glutamate levels with unprecedented spatiotemporal resolution, shedding new light on the dynamics of glutamate neurotransmission in the context of TBI. This represents a step toward better understanding how neurons and astrocytes respond to injury to dynamically alter glutamate signaling and highlights early time windows after TBI as area for therapeutic interventions aimed at reducing pathological glutamate signaling.

## TRANSPARENCY, RIGOR, AND REPRODUCIBILITY

Mice were allocated randomly to distinct experimental cohorts, and all procedures involving CCI, and sham surgeries were conducted on animals aged between 7 to 9 weeks. Sample sizes were determined to guarantee statistical power exceeding 0.8 at a significance level of α=0.05. Automation was employed for ROI mapping when possible, mitigating potential biases from experimenters. Linear mixed modeling was employed, when possible, to accommodate variations both between and within individual animals.

## ACKNOWLEDGMENTS

We thank all the members of the Dulla laboratory for helpful comments about the project and manuscript.

## AUTHORS’ CONTRIBUTIONS

JPG: conceptualization, methodology, software, investigation, writing - original draft. MS: resources, validation, writing – review & editing. ANB: GLAST analysis, writing – review & editing. MA, CD*: conceptualization, methodology, resources, software, writing - original draft.

