## Supplemental Figure 1 for "Glutamate uptake is transiently compromised in the perilesional cortex following controlled cortical impact"

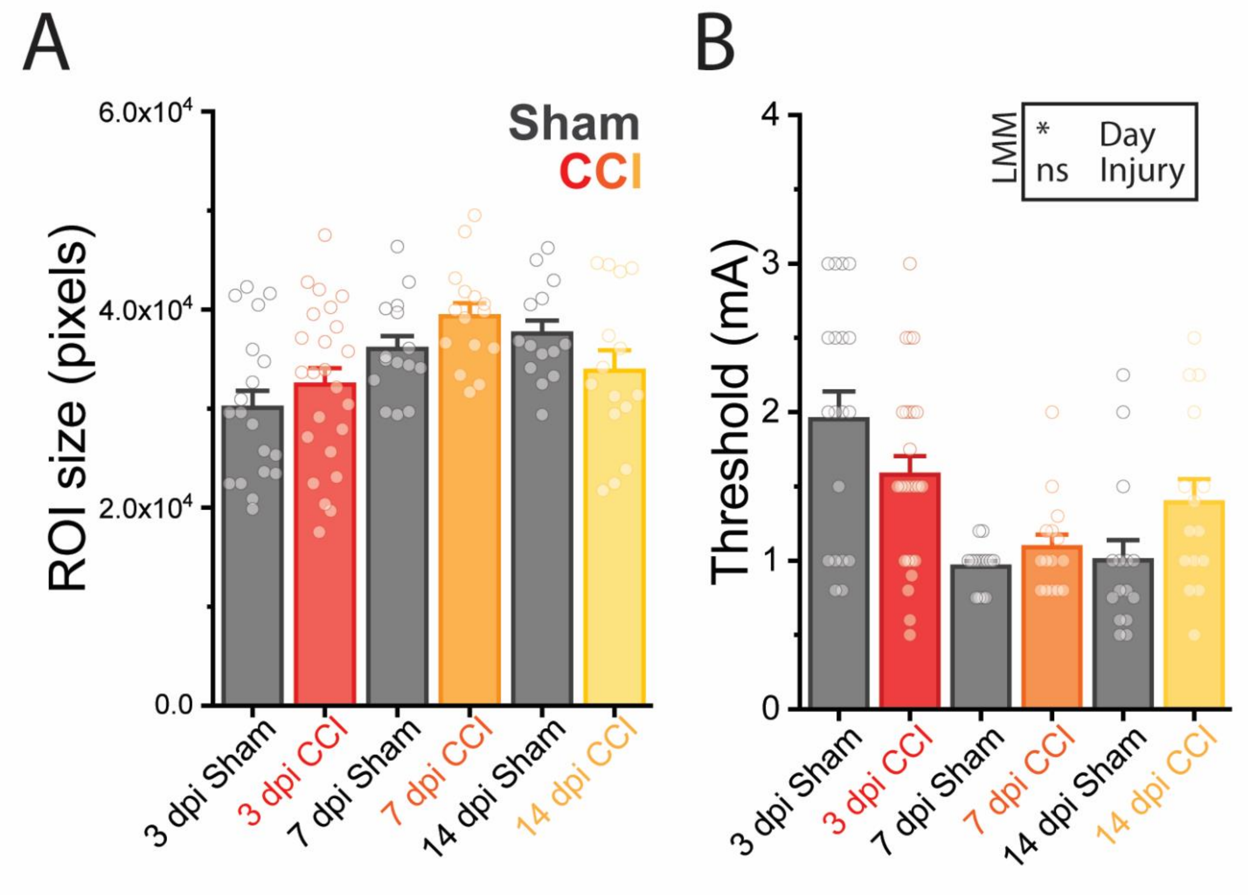


**Supplemental Figure 1. Automated Detection Method for ROI Determination and Threshold Intensity Setting in iGluSnFR Imaging.**

**A.** ROI size was determined using an automated detection method. Pixels that had a peak iGluSnFR signal 2 standard deviations greater than the mean iGluSnFR signal were included in the ROI. There is no significant difference of day or injury. Error bars indicate SEM (3dpi Sham: n= 4 mice, 19 ROIs; 3dpi CCI: n= 5 mice, 25 ROIs; 7dpi Sham: n = 3 mice, 15 ROIs; 7dpi CCI: n = 3 mice, 15 ROIs; 14dpi Sham: n = 3 mice, 15 ROIs; 14dpi CCI: n = 3 mice, 15 ROIs) **B.** Threshold intensity was set at 2x the resolvable threshold stimulation base on iGluSnFR imaging. While days 3 and 7 and 3 and 14 are significantly different ($p = 0.01726$), there is no significant difference of injury at any time point. LMM: p 0.0373, *indicates $P < 0.05$ for differences between Sham and CCI. Error bars indicate SEM (3dpi Sham: n= 4 mice, 19 ROIs; 3dpi CCI: n= 5 mice,

25 ROIs; 7dpi Sham: n = 3 mice, 15 ROIs; 7dpi CCI: n = 3 mice, 15 ROIs; 14dpi Sham: n = 3 mice, 15 ROIs; 14dpi CCI: n = 3 mice, 15 ROIs).
